# Novel tumor suppressor roles for *GZMA* and *RASGRP1* in dissemination of both *Theileria annulata*-transformed macrophages and human B-lymphoma cells

**DOI:** 10.1101/338160

**Authors:** Zineb Rchiad, Malak Haidar, Hifzur Rahman Ansari, Shahin Tajeri, Sara Mfarrej, Fathia Ben Rached, Abhinav Kaushik, Gordon Langsley, Arnab Pain

**Affiliations:** Pathogen Genomics Laboratory, BESE Division, King Abdullah University of Science and Technology (KAUST), Thuwal-23955-6900, Saudi Arabia; Laboratoire de Biologie Cellulaire Comparative des Apicomplexes, Faculté de Médecine, Université Paris Descartes - Sorbonne Paris Cité, France; Inserm U1016, Cnrs UMR8104, Cochin Institute, Paris, 75014 France; King Abdullah International Medical Research Center (KAIMRC), King Abdulaziz Medical City, Ministry of National Guard Health Affairs, Jeddah 21423, Saudi Arabia; Global Station for Zoonosis Control, Global Institution for Collaborative Research and Education (GI-CoRE), Hokkaido University, N20 W10 Kita-ku, Sapporo, Japan

**Keywords:** *Theileria annulata*, *RASGRP1*, *GZMA*, Tumor suppressor, Transcriptome

## Abstract

*Theileria annulata* is a tick-transmitted apicomplexan parasite that infects and transforms bovine leukocytes into disseminating tumors that cause a disease called tropical theileriosis. Using comparative transcriptomics we identified genes transcriptionally perturbed during *Theileria*-induced transformation. Dataset comparisons highlighted a small set of genes associated with *Theileria*-transformed leukocyte dissemination. The roles of Granzyme A (*GZMA*) and RAS guanyl-releasing protein 1 (*RASGRP1*) were verified by CRISPR/Cas9-mediated knock-down. Knocking down of *GZMA* and *RASGRP1* in attenuated macrophages led to a regain in their dissemination in Rag2/γC mice confirming their role as dissemination suppressors *in vivo*. We further evaluated the roles of *GZMA* and *RASGRP1* in human B-lymphoma cells by comparing the transcriptome of 934 human cancer cell lines to that of *Theileria*-transformed bovine host cells. We confirmed dampened dissemination potential of human B-lymphoma cells that overexpress *GZMA* and *RASGRP1*. Our results provide evidence that *GZMA* and *RASGRP1* have a novel tumor suppressor function in both *T. annulata*-infected bovine host cells and in human B-lymphomas.

**Summary:** We compared the transcriptomes of *Theileria annulata* transformed B-lymphocytes to 934 human cancer cell lines and provide functional evidence for shared tumor suppressor roles for GZMA and RASGRP1 in controlling the dissemination phenotype of both human B lymphomas and Theileria-transformed leukocytes.

## INTRODUCTION

*Theileria annulata* is a tick-transmitted apicomplexan parasite that infects and transforms bovine leukocytes into disseminating tumors that cause a widespread disease called tropical theileriosis. In countries endemic for tropical theileriosis live attenuated vaccines are produced by multiples *in vitro* passages of virulent, transformed macrophages and vaccination protects animals from severe disease (1). Amazingly the fully transformed state can be completely reversed by drug-induced parasite death making *Theileria*-infected leukocytes a powerful cellular model to identify genes regulating cellular transformation and dissemination (2). This parasite-based reversible model of leukocyte transformation has allowed the identification of several cell signaling pathways associated with the virulence of *Theileria*-transformed leukocytes such as c-Jun NH2-terminal kinase/c-Jun/PI3 kinase signaling (3), protein kinase A (PKA) (4), transforming growth factor beta 2 (TGF-b2) (5) (6) and SMYD3/MMP9 (7). MMP-9 and c-Jun are associated with invasion, proliferation and angiogenesis in *Theileria*-mediated host cell transformation as well as in human cancer (3, 8-10). Epigenetic changes also contribute to *Theileria*-induced leukocyte transformation (12). OncomiR addiction has been described as being generated by a miR-155 feedback loop in *T. annulata*-transformed B cells (13). Similarly, miR-126-5p contributes to infected macrophage dissemination through JNK-Interacting Protein-2 (JIP2)/JNK1/AP1-mediated *MMP9* transcription (14).

It’s well established that *T. annulata* modulates gene expression of its host cell and hijacks key signalling cascades. For example, RNA extracted from *T. annulata*-transformed B cells was used to screen bovine microarrays demonstrating that infection had reconfigured host cell gene expression (11). Nonetheless, a systematic and genome scale transcriptional comparison of B cells and macrophages transformed by *T. annulata* has been lacking. In this study, we used RNA-seq to define the transcriptional landscapes of two *T. annulata*-transformed B-cell lines and a virulent *T. annulata*-transformed macrophage line (Ode) and the attenuated live vaccine directly derived from it. High stringency bioinformatic comparisons of the transcriptional landscapes identified four candidate genes (*MMP9, GZMA, RASGRP1* and *SEPP1*) as potential players in the dissemination of virulent *T. annulata*-transformed macrophages.

The infection of lymphocytes and macrophages by *Theileria annulata* causes a lymphoproliferative phenotype with properties largely similar to human cancer, most notably immortalization, independence of exogenous growth factors, uncontrolled proliferation and invasiveness. The similarity between *Theileria*-transformed leukocytes and human leukemia suggests that *Theileria-induced transformation* could be a powerful model to elucidate common mechanisms underpinning tumour virulence. In order to generalize our *Theileria*-based observations we compared the transcriptome maps of 934 human cancer cell lines to the transcriptomes of *T. annulata* transformed B-lymphocytes and provide functional evidence for shared tumor suppressor roles for *GZMA* and *RASGRP1* in controlling the dissemination phenotype of both human B lymphomas and *Theileria*-transformed leukocytes.

## Materials and Methods

### Cell lines

The BL3 (15), TBL3, BL20 (16), TBL20 B lymphocytes, and Ode macrophages (17) were cultured in RPMI 1640 medium supplemented with 2 mM of L-glutamine (Lonza, catalogue number 12-702F) and 10 mM Hepes (Lonza, catalogue number 17-737E), 10 % heat-inactivated FBS (Gibco, catalogue number 10082147), 100 units/ml of Penicillin and 100 µg/ml of streptomycin (Lonza, catalogue number 17-602E) and 10mM b-mercaptoethanol (Sigma-Aldrich, catalogue number M6250) for BL3/TBL3 and BL20/TBL20. The virulent (Vir) hyper-disseminating Ode cell line was used at low passage (53-71), while its attenuated (Att) poorly disseminating vaccine counterpart corresponded to passages 309-317. The OCI-LY19 cell line (DSMZ, ACC 528) was cultured in Minimum Essential Medium Eagle - Alpha Modification (Gibco, catalog number 12000063) supplemented with 2.2g/L of sodium bicarbonate (Thermofisher Scientific, catalog number 25080094), 20% FBS, 10 mM Hepes and 100 units/ml of Penicillin and 100 ug/ml of streptomycin. The RI-1 cell line (DSMZ, ACC 585) was cultured in RPMI1640 and supplemented with 10% FBS, 100 units/ml of Penicillin and 100 ug/ml of streptomycin and 10 mM Hepes. All cell lines were incubated at 37°C with 5% CO_2_. All cell lines were regularly tested for mycoplasma contamination.

### RNA extraction

Cells were seeded in 3 biological replicates at a density of 2.5×10^5^ cell/ml. RNA extraction was performed using the PureLink RNA Mini Kit (Life technologies, catalogue number 12183018A) following the manufacturer’s instructions. Briefly, cells were pelleted, lysed and homogenized using a 21-gauge needle, then 70% ethanol was added to the cell lysates and the samples were loaded on spin cartridges to bind RNA. After 3 washes, RNA was eluted in RNase-free water. The quality of extracted RNA was verified using a Bioanalyzer 2100 and quantification carried using Qubit (Invitrogen, catalogue number Q10210).

### Illumina library preparation and sequencing

Strand-specific RNA-sequencing (ssRNA-seq) libraries were prepared using the illumina Truseq Stranded mRNA Sample Preparation Kit (Illumina, catalogue number RS-122-2101) following the manufacturer’s instructions. Briefly, 1ug of total RNA was used to purify mRNA using poly-T oligo-attached magnetic beads. mRNA was then fragmented and cDNA was synthesized using SuperScript III reverse transcriptase (Thermofisher, catalogue number 18080044), followed by adenylation on the 3’ end, barcoding and adapter ligation. The adapter ligated cDNA fragments were then enriched and cleaned with Agencourt Ampure XP beads (Agencourt, catalogue number A63880). Libraries validation was conducted using the 1000 DNA kit on 2100 Bioanalyzer (Agilent Technologies, catalogue number 5067-1504) and quantified using qubit (Thermofisher, catalogue number Q32850). ssRNA libraries were sequenced on Illumina Hiseq2000 and Hiseq4000. The sequenced reads were mapped to the *Bos taurus* genome Btau 4.6.1. The quality of the sequenced libraries is shown in supplementary figure S1.

### Sequencing data analysis

The quality of sequence reads and other parameters were checked using FastQC (http://www.bioinformatics.babraham.ac.uk/projects/fastqc/). The raw RNA-seq reads were processed for adaptor trimming by Trimmomatic (19). The strand-specific reads were mapped on to Bovine genome (bosTau7; Btau_4.6.1; GCF_000003205.5) using Tophat2 (-g 1 --library-type fr-firststrand). The samples with respective replicates were analyzed further for differential gene expression by three different tools, baySeq (20), DESeq2 (21) (fitType =“local”) and CuffDiff2 (22) with default parameters unless mentioned specifically. The count values for DESeq2 and baySeq were calculated from BAM files using HTSeq-count tool (23). The transcriptome quality plots were generated by cummeRbund package (v2.14.0) in R (http://bioconductor.org/packages/release/bioc/html/cummeRbund.html). The sequencing data is available in the NCBI Gene Expression Omnibus, GEO ID: GSE135377.

### Identification of differentially expressed genes after infection and attenuation by comparative transcriptome analysis

The transcriptome data was analyzed with baySeq, DESeq2 and CuffDiff2. A gene was considered as a differentially expressed gene (DEG) if it has a padj<0.05 and a fold change (FC)>2. The final list of DEGs contained genes commonly differentially expressed in CuffDiff2, DESeq2 and baySeq. This approach minimalizes the total number of DEGs for further analysis and allows stringent selection of the most significant and reproducible DEGs.

### qRT-PCR

Total RNA was reverse transcribed using the High Capacity cDNA Reverse Transcription Kit (Applied Biosystems, catalogue number 4368814) as follows: 2 *μ*g of total RNA, 2 *μ*L of RT buffer, 0.8*μ*L of 100mM dNTP mix, 2.0μL of 10X random primers, 1*μ*L of MultiScribe reverse transcriptase and Nuclease-free water to a final volume of 20 *μ*L. The reaction was incubated 10 min at 25°C, 2 h at 37°C then the enzyme inactivated at 85°C for 5 min. Real time PCR was performed in a 10*μ*L reaction containing 20-30 ng cDNA template, 5 *μ*L 2X Fast SYBR Green Master Mix and 500 nM of forward and reverse primers. The reaction was run on the 7500 HT Fast Real-Time PCR System (Applied Biosystems). GAPDH was used as a housekeeping gene and the results were analyzed by the 2^−ΔΔCT^ method. The error bars represent the SEM of 3 biological replicates. Primers were designed and assessed for secondary structures using the Primer Express Software v3.0. The primers of all genes are listed in Table S3.

### Transfection

Macrophages were transfected by electroporation using the Nucleofector system (Amaxa Biosystems). A total of 5 × 10^5^ cells were suspended in 100 *μ*L of Nucleofector V solution mix (Lonza, VCA-1003) with 2 *μ*g of *GZMA* and *RASGRP1* CRISPR/Cas9 plasmids and subjected to nucleofection using the cell line-specific program T-O17. The human B-lymphoma cells were transfected with 2 *μ* g of *GZMA* (SCBT, sc-403958-ACT), *RASGRP1* (SCBT, sc-402120-ACT) and *SEPP1* (SCBT, sc-417457-ACT) activation plasmids, and the non-specific control plasmid (SCBT, sc-437275) using the T-O17 program. After transfection, cells were suspended in fresh complete medium and incubated at 37°C with 5% CO_2_.

### Matrigel chamber assay

The invasive capacity of Ode macrophages was assessed *in vitro* using matrigel migration chambers, as described in (3). The CultureCoat Medium basement membrane extract (BME) 96-wells cell invasion assay was performed according to Culturex instructions (Trevigen, catalog number 3482-096-K). After 24 h of incubation at 37°C, each well of the top chamber was washed once in buffer. The top chamber was placed back onto the receiver plate. One hundred microliters of cell dissociation solution-Calcein AM was added to the bottom chamber of each well, and the mixtures were incubated at 37°C for 1 h with fluorescently labeled cells to dissociate the cells from the membrane before reading at 485-nm excitation and 520-nm emission wavelengths.

### Soft agar colony forming assay

A two-layer soft agar culture system was used. Cell counts were performed by ImageJ software. A total of 2,500 cells were plated in a volume of 1.5 ml (0.7% bacto Agar+2× RPMI 20% Fetal bovine Serum) over 1.5 ml base layer (1% bacto agar +2× RPMI 20% Fetal bovine Serum) in 6-well plates. Cultures were incubated in humidified 37°C incubators with an atmosphere of 5% CO_2_ in air, and control plates were monitored for growth using a microscope. At the time of maximum colony formation (10 days in culture), final colony numbers were counted after fixation with 0.005% Cristal Violet.

### Intracellular levels of hydrogen peroxide (H_2_O_2_)

Cells were seeded at 1×10^5^ cell/well in a 96 well plate and incubated in complete medium for 18 h prior to the assay. Cells were then washed with PBS and incubated with 100 *μ*L of 5 M H2-DCFDA in PBS (Molecular Probes, catalogue number D399). H_2_O_2_ levels were assayed on a fusion spectrofluorimeter (PackardBell) by spectrofluorimetry at 485 and 530nm excitation and emission wavelengths respectively.

### *In vivo* mouse studies and quantification of *Theileria annulata*-transformed macrophages load in mouse tissues

*T. annulata*-infected macrophage cell lines (Virulent Ode passage 53, attenuated Ode passage 309, attenuated Ode transfected with *RASGRP1* CRISPR/Cas9 knock out plasmid and attenuated Ode transfected with *GZMA* CRISPR/Cas9 knock out plasmid) were injected into four groups of five Rag2*γ*C immunodeficient mice that were equally distributed on the basis of age and sex in each group. The injection site was disinfected with ethanol and one million cells (in 200 µl PBS) were injected under the skin after gentle shaking of the insulin syringe. The mice were kept for 3 weeks and then they were humanely sacrificed and dissected. Six internal organs including heart, lung, spleen, mesentery, left kidney and liver were taken and stored in 500 *µ* L PBS in Eppendorf tubes at −20°C. The tissues were subjected to genomic DNA extraction using the QIAmp DNA mini kit (Qiagen, catalogue number 51304). DNA concentrations were measured by Nanodrop™ 1000 spectrophotometer (Thermo Fischer scientific) and before each quantitative PCR reaction samples were diluted to give a DNA concentration of 0.5 ng/*µ* L. Absolute copy numbers of a single copy *T. annulata* gene (*ama-1*, TA02980) that is representative of *T. annulata*-infected macrophage load in each tissue were estimated by the method described in (24), with some modifications. *ama-1* was cloned into pJET 1.2/blunt cloning vector using CloneJET PCR Cloning Kit (Thermo scientific, catalogue number K1232). The cloned plasmid was amplified in DH5-Alpha cells and purified with QIAfilter™ Plasmid Maxi Kit (Qiagen, catalogue number 12243). Plasmid concentration was measured using Qubit (Thermofisher, catalogue number Q32850). The primers for cloning were: forward 5’-GGAGCTAACTCTGACCCTTCG-3’and reverse 5’-CCAAAGTAGGCCAATACGGC-3’. Quantitative PCR primers were: forward 5’-GACCGATTTCATGGCAAAGT-3’and reverse 5’-TTGGGGTCATGATGGGTTAT-3’.

### Transcriptome-based clustering of *Theileria*-transformed bovine host cells and human cancer cell lines

The processed and quality trimmed reads from TBL20/BL20 and BL3/TBL3 samples were mapped to *Bos taurus* UMD3.1 genome using HISAT2 software (25) with default settings. The mapped reads were used for gene-level TPM quantification using StringTie (Version 1.3.3b) (26, 27). The quantified genes were converted into their human ortholog Ensemble gene ID by finding one-to-one orthologs between human and *B. taurus* genomes using OMA browser (28). Subsequently, TPM values of transcripts expressed across 934 human cancer cell lines were obtained from EBI cancer cell line Expression Atlas (29). The redundant transcripts in the cancer cell line expression set were collapsed using collapseRow function from the WGCNA R package (30). Using the common human ensemble gene ID, gene expression matrices of *B. taurus* and human cell lines were merged together, which was then subjected to hierarchical clustering of samples using HCPC (31) with 3 Principal Components (nPCs), which resulted in 4 broad clusters. The sub-cluster containing the human cancer cell lines along with TBL20/BL20 and BL3/TBL3 were shortlisted for further analysis. Whereby, samples were scanned for similar gene expression profiles by computing the adjacency of each shortlisted sample with the rest of the samples, using adjacency function (method=“Distance”) from WGCNA R package. The resultant adjacency matrix was then subjected to flashClust (31) program for computing the dendrogram for manual inspection of TBL20/BL20 and BL3/TBL3 containing sub-cluster. A schematic of the used pipeline is presented in Fig. S6. The complete dendrogram of the 934 human cancer cell lines and *Theileria*-tranformed lymphocytes can be viewed in Fig. S7.

### Ethics statement

The protocol (12-26) was approved by the ethics committee for animal experimentation of the University of Paris-Descartes (CEEA34.GL.03312). The university ethics committee is registered with the French National Ethics Committee for Animal Experimentation that itself is registered with the European Ethics Committee for Animal Experimentation. The right to perform the mice experiments was obtained from the French National Service for the Protection of Animal Health and satisfied the animal welfare conditions defined by laws (R214-87 to R214-122 and R215-10) and GL was responsible for all animal experimentation, as he holds the French National Animal Experimentation permit with the authorisation number (B-75-1249). This project is also covered by the KAUST IBEC number 19IBEC12.

## RESULTS

### Differentially expressed bovine genes in *T. annulata*-transformed leukocytes

The infection and full transformation of the BL20 cell line with *T. annulata* caused profound transcriptional changes, as previously reported for infected BL20 cells (11). Similarly, infection of BL3 cell with *T. annulata* also provoked changes in host cell gene expression (Fig. S2a). Transcriptional changes between virulent compared to attenuated Ode macrophages are less profound, likely because the macrophages only appear to differ in dissemination potential (Fig. S2a).

To identify bovine genes whose transcription is perturbed by transformation and attenuation of dissemination of *T. annulata*-transformed leukocytes we concentrated on the most differentially expressed genes (DEGs) (Fig. 1, Table S1). Many of these genes are annotated as being implicated in cell proliferation and metastasis. Amongst the top five-upregulated transcripts in TBL20 is *MMP9* (matrix metallopeptidase 9), a gene highly expressed in different cancer types and linked to metastasis and angiogenesis (32). *WC1-8* is the third most upregulated gene in TBL20 lymphocytes and has been described as being also upregulated in ovarian carcinoma cells (33). The most down-regulated transcripts in TBL20 cells include *LAIR1* (leukocyte associated immunoglobulin like receptor 1) and *VPREB* (pre-B lymphocyte 1). *LAIR1* is a strong inhibitor of natural killer cell-mediated cytotoxicity and an inhibitory receptor, which down-regulates B lymphocyte immunoglobulin and cytokine production (34). Down-regulation of *LAIR-1* was not unexpected, as it’s loss of expression is observed during B cell proliferation (35). *ZBTB32* (zinc finger and BTB domain containing 32), *IL21R* (interleukin 21 receptor), and *MMP9* are also among the top five up-regulated transcripts in infected TBL3 B lymphocytes. The five most down-regulated transcripts in TBL3 are *KRT6C* (keratin 6C), *MATK* (megakaryocyte-associated tyrosine kinase), *IGSF9B* (immunoglobulin superfamily member 9B), *A2M* (alpha-2-macroglobulin) and *H2AFY2* (H2A histone family, member Y2). The biological functions of these genes include inhibition of cell growth and proliferation (36), repression of DNA transcription (37) and inhibition of cell adhesion and migration (38), functions that are often dampened to allow continuous proliferation and survival of transformed cells. We confirmed by qRT-PCR differential expression of 21 randomly selected genes from the BL20/TBL20 and BL3/TBL3 RNA-seq datasets (Fig. S2b).

**Figure 1:**
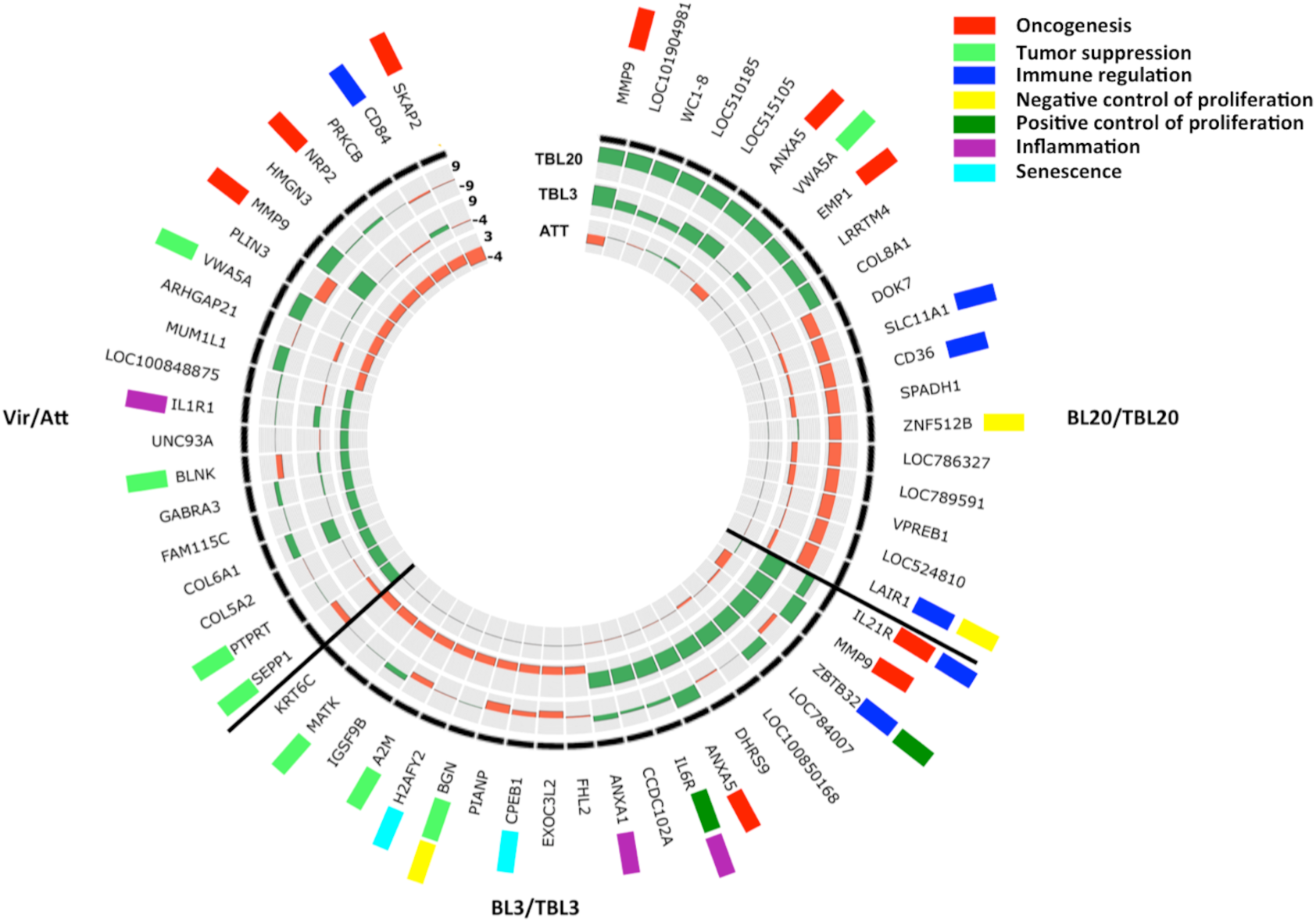
Top 20 differentially expressed genes in Theileria-transformed bovine host cells. **A** Circos plot showing the top 10 up- and down-regulated DEGs in BL3/TBL3, BL20/TBL20 and Attenuated versus Virulent (Att/Vir) Ode macrophages. The circular heatmap represents the FC of the top DE genes in BL20/TBL20, BL3/TBL3, and Att/Vir Ode in the outer, middle and inner rings, respectively, where green reflects the level of up-regulation and red down-regulation. The genes with biological functions related to tumorigenesis and immune regulation are tagged with colored rectangles. Genes with no tag are hypothetical genes or have no known function in tumorigenesis and immune regulation.

### Identification of key genes potentially involved in *Theileria*-mediated macrophage dissemination

The most down-regulated transcripts in attenuated Ode macrophages are *SKAP2* (src kinase associated phosphoprotein 2), a gene known to promote tumor metastasis through the regulation of podosome formation in macrophages (39) and *NRP2* that regulates tumor progression by promoting TGF-β-signaling (40). Down-regulation of these genes correlates with decreased dissemination of attenuated macrophages, as previously we have described loss of TGF-β-signaling as being associated with decreased dissemination (5). By contrast, the most highly upregulated transcripts include *SEPP1* (selenoprotein P) and *PTPRT* (protein tyrosine phosphatase, receptor type T). Both *SEPP1* and *PTPRT* have been previously described as a tumor suppressor gene (41, 42). Taken together, the identity of the most strongly up- and down-regulated genes argues that our differential transcription screen could identify novel genes regulating *Theileria*-transformed macrophage dissemination.

To define the genes likely playing important roles in transformation and dissemination, we compared genes differentially expressed (DE) in TBL3, TBL20 and attenuated Ode macrophages. We assumed that the genes most likely to play a key role are upregulated after infection and downregulated upon attenuation, and vice versa. This approach identified four genes likely to play key roles in the dissemination of *Theileria*-transformed leukocytes (Fig. 2a).

**Figure 2:**
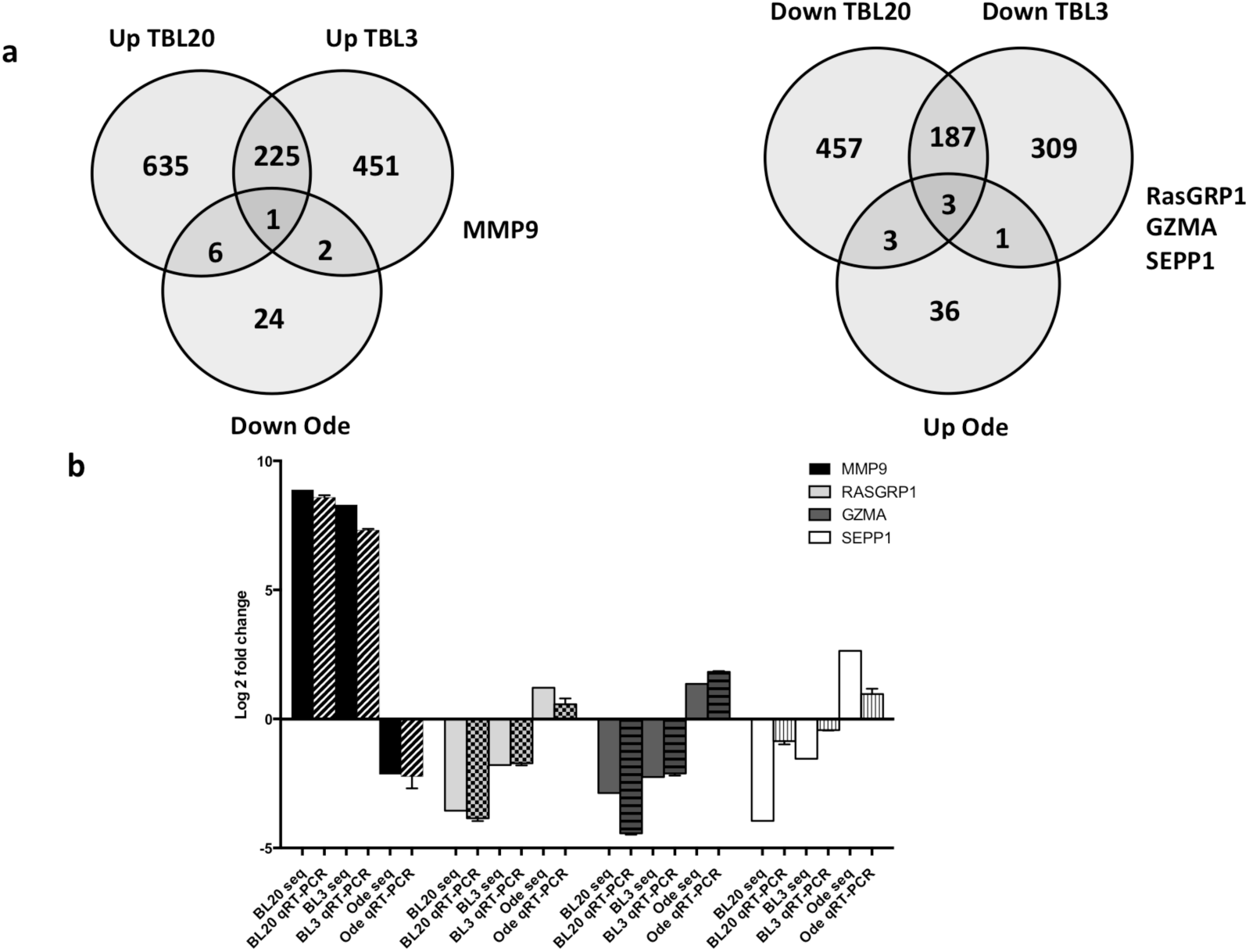
Inversely DEGs in TBL20, TBL3 and Att Ode leukocytes. (a) Venn diagrams illustrating the genes inversely DE in TBL3, TBL20 and Attenuated Ode macrophages. (b) qRT-PCR confirmation of DEGs potentially playing key roles in leukocyte transformation and dissemination. The reactions were set in 3 biological replicates and the fold-change calculated with the 2^Δ Δ ct^ method. The error bars represent SEM.

The genes are *MMP9, SEPP1, GZMA* and *RASGRP1* and their biological functions have been implicated in metastasis and cell invasion (32), selenium transport (43), peptide cleavage by immune cells (44) and regulation of B cell-development and homeostasis and differentiation (45), respectively (Table 1). Differential expression of these genes was confirmed by qRT-PCR (Fig. 2b). We focused on *GZMA, RASGRP1* and *SEPP1*, as the role of *MMP9* in metastasis/dissemination is well established including in *Theileria*-transformed macrophages (46). CRISPR/Cas9-mediated loss of *SEPP1* in attenuated macrophages resulted in a lethal phenotype highlighting its essentiality in transformed macrophage survival and the role of SEPP1 in *Theileria*-transformed cell lines was not further evaluated.

**Table 1:** Biological functions of DEGs potentially playing key roles in *T. annulata*-mediated leukocyte transformation and dissemination

| Gene symbol | Log2 FC (TBL20) | Adj p value | Log2 FC (TBL3) | Adj p value | Log2 FC (ATT) | Adj p value | Biological functions | References on biological functions |
| --- | --- | --- | --- | --- | --- | --- | --- | --- |
| *MMP9* | 8.87 | 0 | 8.29 | 0 | -2.13 | 5.14 E-94 | Metastasis formation, cancer cells invasion | (32) |
| *SEPP1* | -3.95 | 8.74 E-174 | -1.54 | 3.75 E-248 | 2.63 | 1.01 E-140 | Transports selenoprotein, tumor suppressor | (48), (42), (61) |
| *GZMA* | -2.87 | 1.45 E-05 | -2.25 | 1.97 E-41 | 1.02 | 9.14 E-21 | Plays a role in killing pathogen infected cells and cancer cells, induces increase in ROS | (44), (51) |
| *RASGRP1* | -3.55 | 0 | -1.78 | 1.97 E-41 | 1.2 | 2.15 E-15 | Required for correct functioning of lymphocytes in chronic infections, | (45) |

### Ablation of *GZMA* and *RASGRP1* by CRISPR/Cas9 knockdown

To confirm a role for *GZMA* and *RASGRP1* in dissemination of *T. annulata*-transformed Ode macrophages, we knocked down their expression by CRISPR/Cas9 and decreased expression led to a regain in dissemination potential, as estimated in matrigel traversal assays (Fig. 3b). Both GZMA and RASGRP1 have therefore, the potential to function as suppressors of tumor dissemination and consistently, knockdown of *GZMA* also led to a regain in the ability of attenuated macrophages to form colonies in soft agar (Fig.3c). Taken together it strongly suggests that GZMA and RASGRP1 function as tumor suppressors.

**Figure 3:**
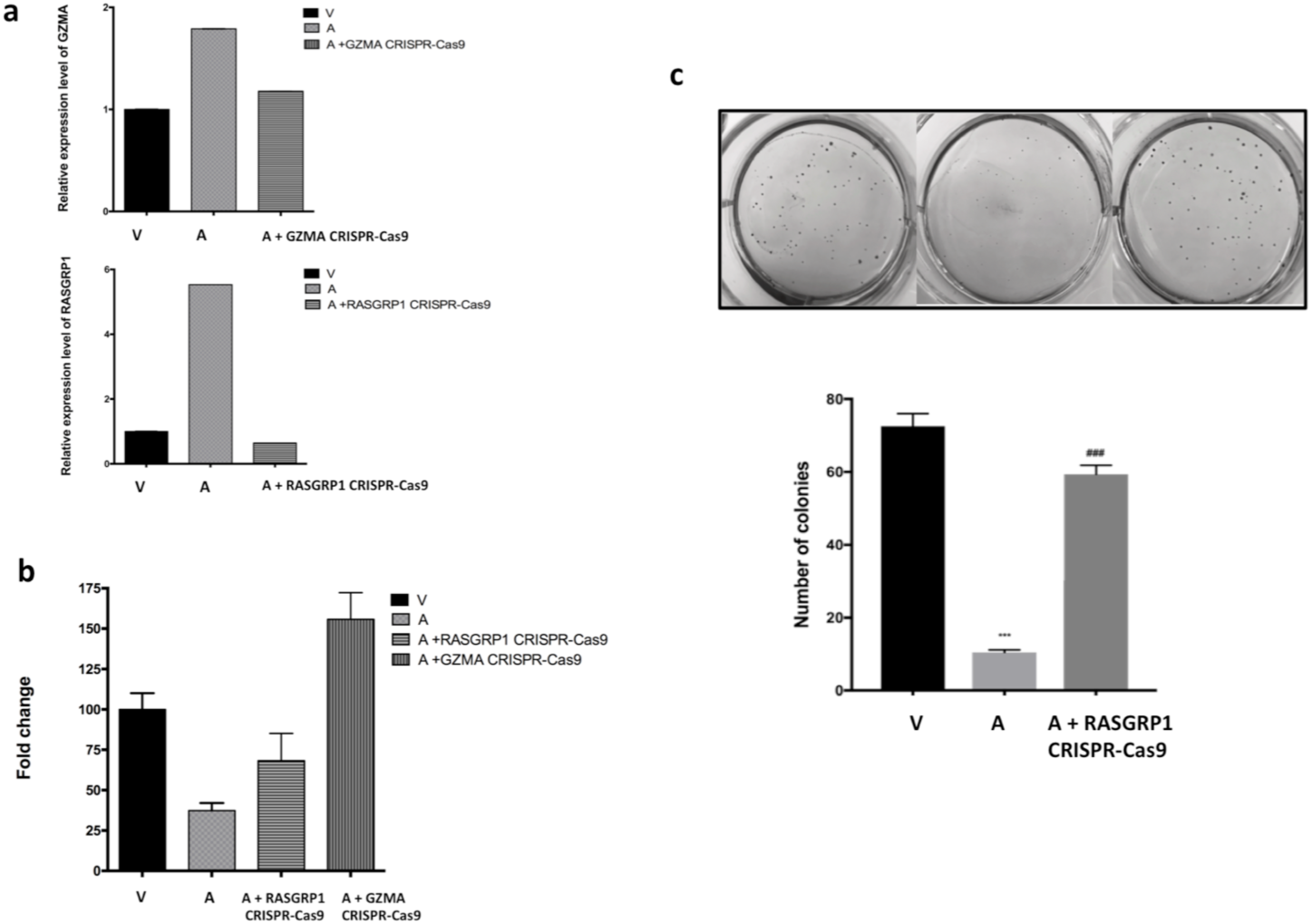
Colony formation on soft agar. (a) qRT-PCR confirmation of *GZMA* (top panel) and *RASGRP1* (bottom panel) knockdown. b) Matrigel chamber assay showing a regain in Matrigel traversal after *RASGRP1* and *GZMA* knockdown. c) Increased colony formation in soft agar following *RASGRP1* knockdown. Non-transfected virulent disseminating Ode macrophages are indicated by V, and non-transfected poorly disseminating attenuated Ode macrophages by A. Error bars represent SD of 3 biological replicates. *** and ### represent student t test p<0.001 compared to virulent and attenuated Ode macrophages, respectively.

### *GZMA* and *RASGRP1* dampen *in vivo* dissemination of Ode macrophages

Similar to metastatic tumor cells *T. annulata*-transformed leukocytes also disseminate in immuno-deficient mice to distant organs and form proliferative foci (47). Dissemination of *Theileria*-transformed leukocytes has been previously attributed to increased production of matrix metaloproteinases (MMPs) (46). As *GZMA* and *RASGRP1* knockdown led to a regain in matrigel traversal we used Rag2*γ*C immunodeficient mice to test for a regain in dissemination *in vivo*. The CRISPR/Cas9-induced ablation of expression of GZMA and RASGRP1 gave rise to an increase in the number of *Theileria*-containing tumors in heart, lung and mesentery, while knockdown of *RASGRP1* increased the number of tumors in the liver (Fig. 4). Thus, loss of *RASGRP1* and *GZMA* expression led to a regain in the invasive capacity of *T. annulata*-transformed macrophages into these organs.

**Figure 4:**
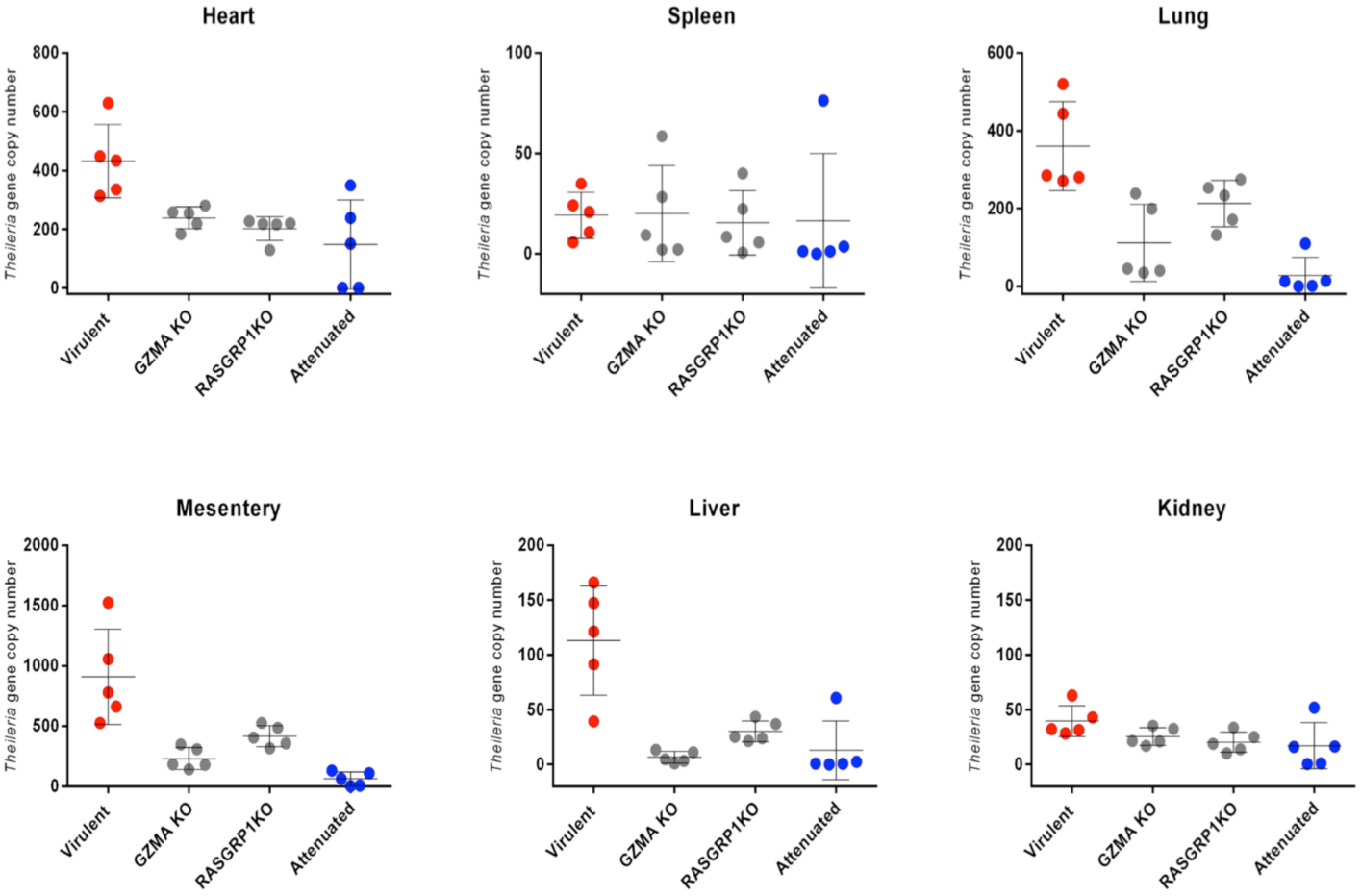
Effect of *GZMA* and *RASGRP1* knockdown on transformed macrophage dissemination *in vivo*. Panels represent the copy number of the single copy *T. annulata* gene (*ama-1*, TA02980) in six internal organs: heart, lung, spleen, mesentery, left kidney and liver. Transformed macrophages were injected into five Rag2*γ*C immunodeficient mice and plotted values represent the mean of obtained *T. annulata-*specific *ama1* gene copy number. Error bars represent SD of 5 biological replicates.

### Induced expression of *GZMA, RASGRP1* and *SEPP1* reduces human B-lymphoma cell dissemination

In order to extend the roles of GZMA, RASGRP1 and SEPP1 to human cancer we sought human tumor cells displaying transcriptional signatures similar to *T. annulata*-transformed leukocytes. To this end, the transcriptional profiles of 934 human cancer cells were obtained from the EBI cancer cell line expression atlas (29) and their profiles compared to those of *Theileria*-transformed TBL3 and TBL20 B-lymphocytes (Fig. 5a). Among the subset human B lymphomas OCI-LY19 and RI-1 displayed the transcriptional signature most similar to TBL3 and TBL20 and therefore, were used to test if *GZMA, RASGRP1* and *SEPP1* can act as suppressors of dissemination in certain types of human cancer. CRISPR-mediated transcriptional activation of *GZMA, RASGRP1* and *SEPP1* resulted in decreased matrigel traversal of the OCI-LY19 B lymphoma. By contrast, only upregulation of *GZMA* showed a statistically significant decrease in traversal of the RI-1 B lymphoma (Fig. 5c). Clearly, *GZMA* has the potential to act as a suppressor in two independent human B-lymphomas, whereas *RASGRP1* and *SEPP1* may only function as suppressors in specific B-lymphomas.

**Figure 5:**
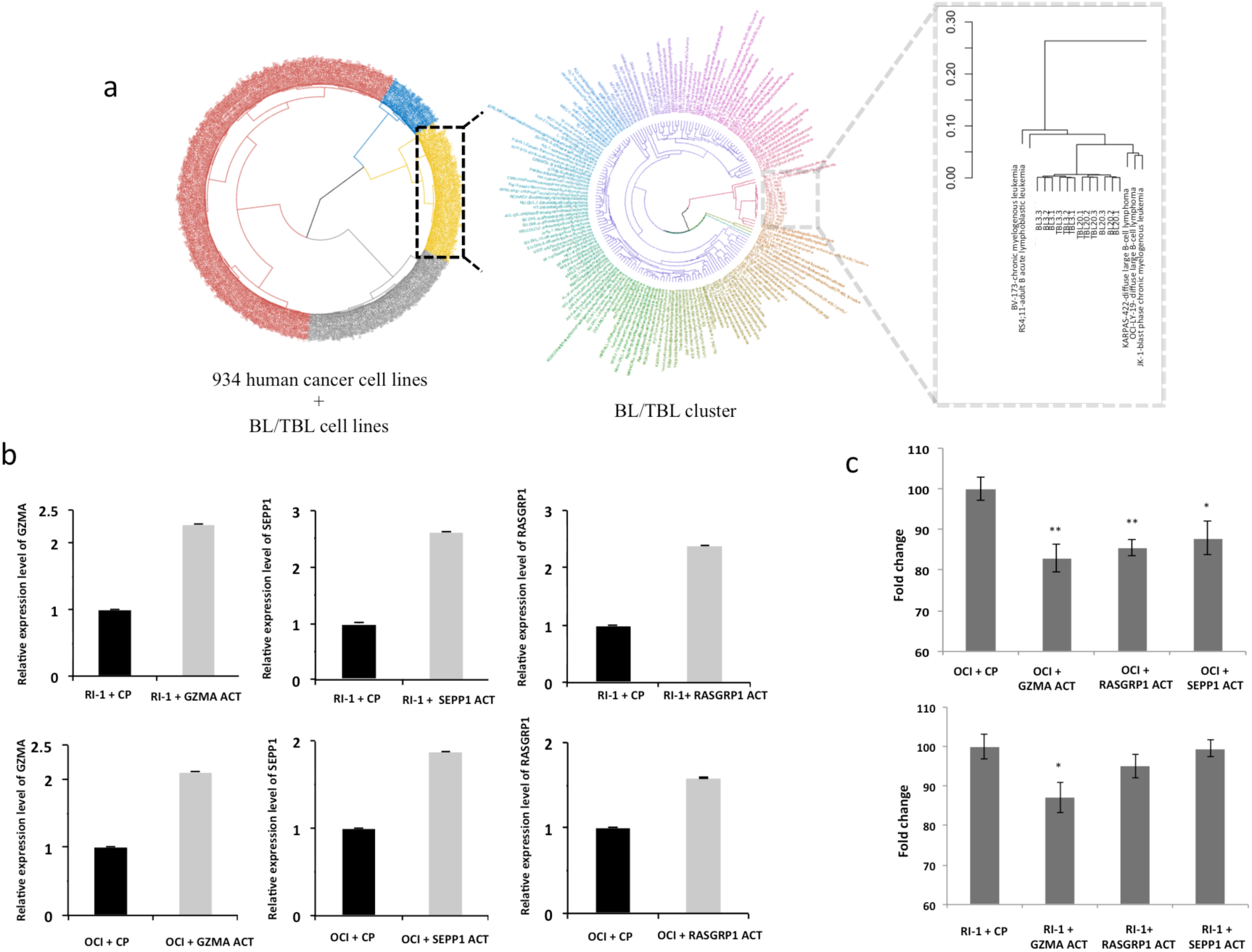
Effect of GZMA, RASGRP1 and SEPP1 activation on human B-lymphomas. (a) (Left panel) PCA based hierarchical clustering of 934 human cancer cell lines and *T. annulata-*transformed bovine B cells and their non-infected counterparts. The cluster containing bovine cells mainly contains leukemic human cancer cell lines. The samples are colored by cluster ID. (Middle cluster) The samples from the BL3/TBL3 and BL20/TBL20 sub-cluster were re-clustered by comparing the similarity of their gene expression profiles. The sample labels are colored by their similarity to each other. (Right panel) The sub-cluster containing bovine cell lines and human cancer cell lines with similar gene expression profile. (b) qRT-PCR determination of *GZMA, RASGRP1* and *SEPP1* expression after CRISPR-mediated gene activation in RI-1 (top panel) and OCI-LY19 (bottom panel). The error bars represent SEM of 3 biological replicates. (c) Matrigel chamber assay illustrating the dissemination potential of RI-1 and OCI-LY19 following activation of *RASGRP1* and *GZMA* transcription. Errors bars represent SEM values of 3 biological replicates * and ** represent student t test p<0.05 and p<0.005, respectively.

## DISCUSSION

In this study, we provide a holistic view of the transcriptional landscape of two *T. annulata*-transformed B cell lines, TBL3 and TBL20, and in addition the landscape of virulent versus attenuated Ode macrophages. In order to find genes with commonly perturbed transcription the different datasets were compared using three independent pipelines to identify just four genes, as potential regulators of tumor dissemination. In addition to *MMP9* three other genes (*SEPP1, GZMA* and *RASGRP1*) were identified as potentially having a role in dissemination. SEPP1 is a major selenoprotein involved in selenium transport and cellular defense against oxidative stress (48). Attenuated macrophages did not survive CRISPR/Cas9-knockdown of *SEPP1* implying death might be due to a failure to control excessive oxidative stress, since attenuated macrophages display high levels of H_2_O_2_ (49).

RASGRP1-activated Ras family proteins possess both pro- and anti-oncogenic properties, depending on the downstream effector pathway and cellular context; reviewed in (50). Our transcription profiling showed that most of the members of the *RASGRP* gene family (*RASGRP1, 2 & 4*) are significantly downregulated in TBL3 and TBL20 (Fig. S3). GZMA is a serine protease that contributes to killing of both tumors and pathogen-infected cells via a caspase-independent pathway (44). *GZMA* expression induces reactive oxygen species (51) and attenuated macrophages are known to be more oxidatively stressed than virulent macrophages (49). Indeed, H_2_O_2_ output was reduced in attenuated macrophages following CRISPR/Cas9-mediated *GZMA* knockdown (Fig. S4). Furthermore, it has been shown that *RASGRP1*-deficient CD8 T cells exhibit markedly reduced expression of GZMB (52). This led us to investigate whether loss of *RASGRP1* could perhaps provoke a drop in *GZMA* expression rendering attenuated macrophages doubly deficient in dissemination *in vivo*. As hypothesized, we found that expression of *GZMA* decreased after *RASGRP1* knockdown (Fig. S5). Moreover, *GZMA* and *RASGRP1* expression is repressed by TGF-β (52, 53) and the role of TGF-β in regulating dissemination of *Theileria*-transformed macrophages is very well established (4-6). Expression of *GZMA* and *RASGRP1* was decreased in attenuated Ode macrophages treated with TGF-β (Table S4). Taken altogether, it suggests that one way TGF-β promotes dissemination of *Theileria*-transformed leukocytes could be via repression of both *GZMA* and *RASGRP1* transcription and their impact on dissemination confirmed *in vivo* in mice. In addition to the role of *GZMA, RASGRP1* and *SEPP1* in regulating tumorigenesis of *Theileria-*transformed leukocytes, all three genes also dampened the capacity of the human OCI-LY19 B-lymphoma to traverse matrigel and GZMA also dampened traversal of human RI-1 B lymphoma cells.

GZMA is known to cleave APEX1 (apurinic/apyrimidinic endodeoxyribonuclease 1) after Lys31 and destroys its oxidative repair functions. APEX1 is involved in NK-cell-mediated killing via GZMA (51, 54) and can suppress the activation of PARP1 during repair of oxidative DNA damage (55), and prevent oxidative stress by negatively regulating Rac1/GTPase activity (56). Additionally, APEX1 directly reduces the redox-sensitive cysteine residues of target transcription factors, enhancing their DNA binding and transcriptional activity. Analysis of our deep RNA-seq data revealed that *APEX1* is downregulated in attenuated macrophages and its expression increases after TGF-β treatment, along with an important downregulation of *GZMA* and *RASGRP1* (Table S4). It is over-expressed in many cancers (57) (58) (59) and has been implicated in growth, migration and invasion of colon cancer both *in vitro* and *in vivo* (60). Interestingly, APEX-1 protects melanoma cells from H_2_O_2_ induced apoptosis (57). The established function of APEX1 in human cancer sustains our novel observations on the tumor suppressor roles of *GZMA* and *RASGRP1*. The ensemble of our results allows us to propose a model, where TGF-β modulates tumor redox balance-mediated dissemination by regulating a GZMA/RASGRP1/APEX1 pathway (Fig. 6).

**Figure 6:**
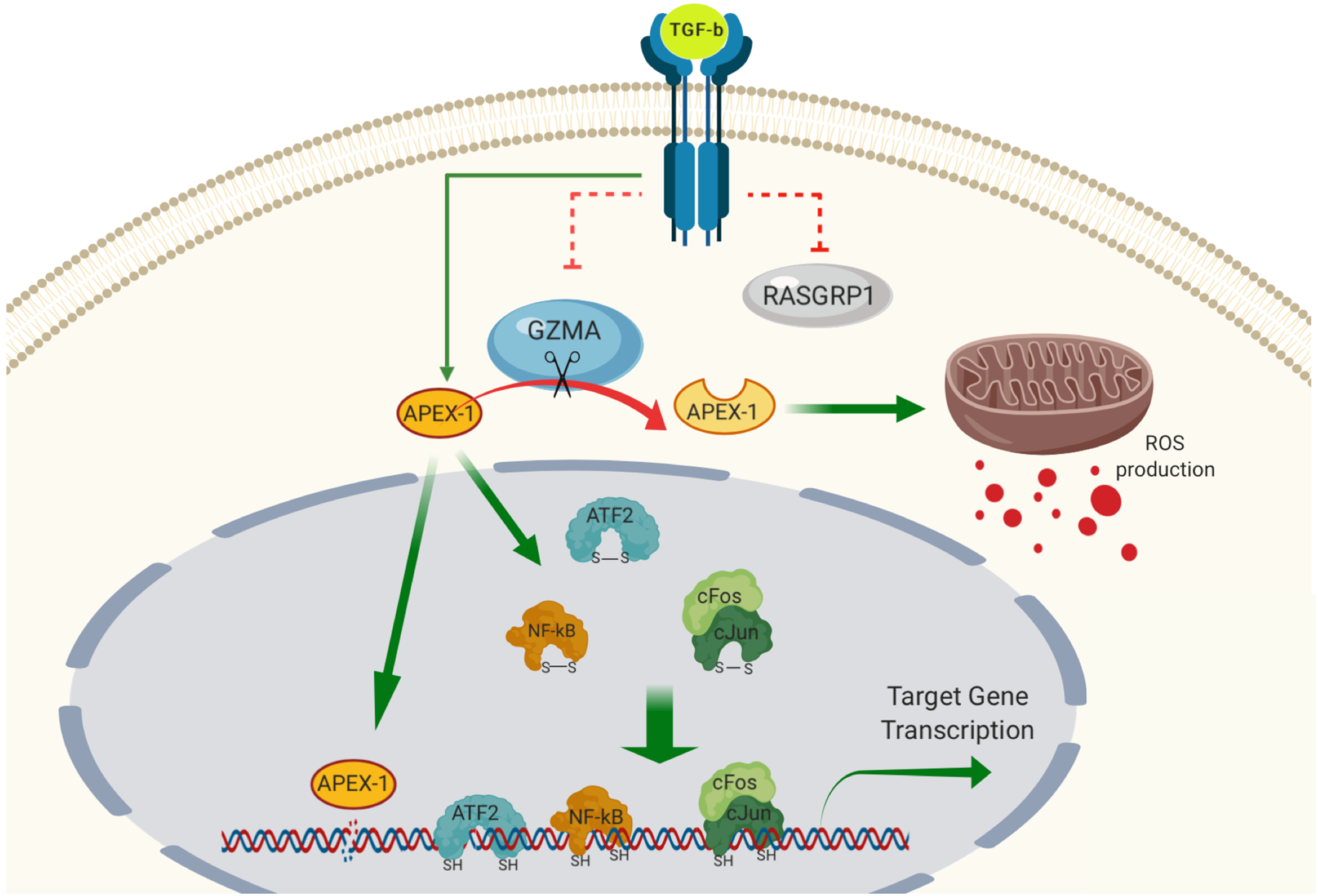
Scheme of proposed TGF/GZMA/RASGRP1/APEX1 pathway. In virulent hyper-disseminating macrophages enhanced TGF-β2 signaling results in upregulated *APEX1* transcription and suppressed *GZMA/RASGRP1* transcription. By contrast, in macrophages attenuated for dissemination reduced TGF-β2 signaling leads to lower *APEX1* transcription and increased in *GZMA* expression. Higher levels of GZMA cleave lower levels of APEX-1 ablating its oxidative repair functions leading to increased ROS output that typifies attenuated macrophages. In virulent, macrophages TGF-β2-induced APEX-1 reduces oxidized forms of target transcription factors such as NF-κB, ATF2, cFos and cJun to increase their DNA binding activity that augments transcription of their target genes to promote cellular transformation and tumor dissemination.

This study has revealed new players in dissemination and oxidative stress regulation of *Theileria*-transformed leukocytes and has provided evidence for similar roles for GZMA and RASGRP1 in transcriptionally matched human B-lymphoma cell lines. The similarity between *Theileria*-induced B cell transformation and human B-lymphomas has unveiled novel therapeutic targets to treat cancer.

## Acknowledgements

This study was supported by a Competitive Research Grant from the Office for Sponsored Research (OSR-2015-CRG4-2610) at King Abdullah University of Science and Technology (KAUST) awarded to AP and GL. ZR acknowledges KAUST for awarding her PhD studentship. ST was supported by a post-doctoral fellowship from ParaFrap (ANR-11-LABX-0024) and in addition to ANR-11-LABX-0024 GL also acknowledges core support from INSERM and the CNRS. We thank members of the Bioscience Core Laboratory (BCL) at KAUST for producing the raw sequencing datasets. Franck Letourneur of the genomics platform at the Cochin institute (GENOM’IC) for quantifying the pJET-*ama-1* plasmid.

## Author contributions

AP and GL conceived and designed the study. ZR prepared the ssRNAseq libraries, SM, MH and ZR ran qRT-PCR reactions. MH performed the soft agar colony formation assay, intracellular H_2_O_2_ levels and invasion assays. ST performed the mouse dissemination assays, HRA, AK and ZR performed data analysis. ZR and MH prepared the figures and ZR prepared the first draft of the manuscript with input from FBR that was then edited by MH, GL and AP.

## Supplementary figure legends

**Figure S1: Sequencing quality of all samples.**

(a) Clustering of all samples. (b) Density plot representing FPKM distribution of all samples.

**Figure S2: Differentially expressed genes in TBL20, TBL3 and Attenuated Ode leukocytes.**

(a) Histogram showing the number of up- (dark grey) and down- (light grey) regulated genes in all 3 datasets. The area-proportional Venn diagrams represent the intersection between the lists of DEGs from CuffDiff2 (green), DESeq2 (black) and baySeq (blue). The intersection between the 3 pipelines reflects the number of up- and down-regulated genes. The list of DEGs is listed in Table S2. (b) qRT-PCR confirmation of randomly selected genes in TBL20 and TBL3. The reactions were set in 3 biological replicates and the fold change calculating with the 2^ΔΔct^ method. The error bars represent SEM.

**Figure S3: Log2FC values of RASGRP1-4 in TBL20 and TBL3**

RNAseq Log2FC values from DESeq2 of TBL20 and TBL3 compared to BL20 and BL3, respectively.

**Figure S4: Effect of *GZMA* knockdown on H**_**2**_**O**_**2**_ **output.**

H_2_O_2_ output by virulent (V), attenuated (A), and attenuated Ode macrophages after CRISPR/Cas9-mediated *GZMA* knockdown. Error bars represent SD of 3 biological replicates. ** and ## represent p<0.01 compared to virulent and attenuated Ode macrophages, respectively.

**Figure S5: Effect of *RASGRP1* knockdown on *GZMA* expression.**

qRT-PCR of GZMA in virulent (V), attenuated (A) and attenuated Ode macrophages after CRISPR/Cas9-mediated *RASGRP1* knockdown. Error bars represent SD of 3 biological replicates *** and ### represent p<0.001 compared to virulent and attenuated Ode macrophages, respectively.

**Figure S6: Schematic of *T. annulata* host cells and human cancer cell lines transcriptome clustering**

**Figure S7: Complete cluster of 934 human cancer cell lines and BL20/TBL20, BL3/TBL3**

**Table S1: Top 5 up- and down-regulated DEGs in infected and attenuated cell lines.**

This list was generated based on FC values of DEGs listed in Table S2.

**Table S2: List of DEGs in infected and attenuated cell lines.**

For details on the methods used to generate this list, please see in the materials and methods section.

**Table S3: List of qRT-PCR primers.**

**Table S4: List of DEGs in attenuated Ode macrophages after TGF-β2 treatment.**

Genes listed as DE from DESeq2

