## Supplementary figures and images for "Novel tumor suppressor roles for *GZMA* and *RASGRP1* in dissemination of both *Theileria annulata*-transformed macrophages and human B-lymphoma cells"

### Fig S1

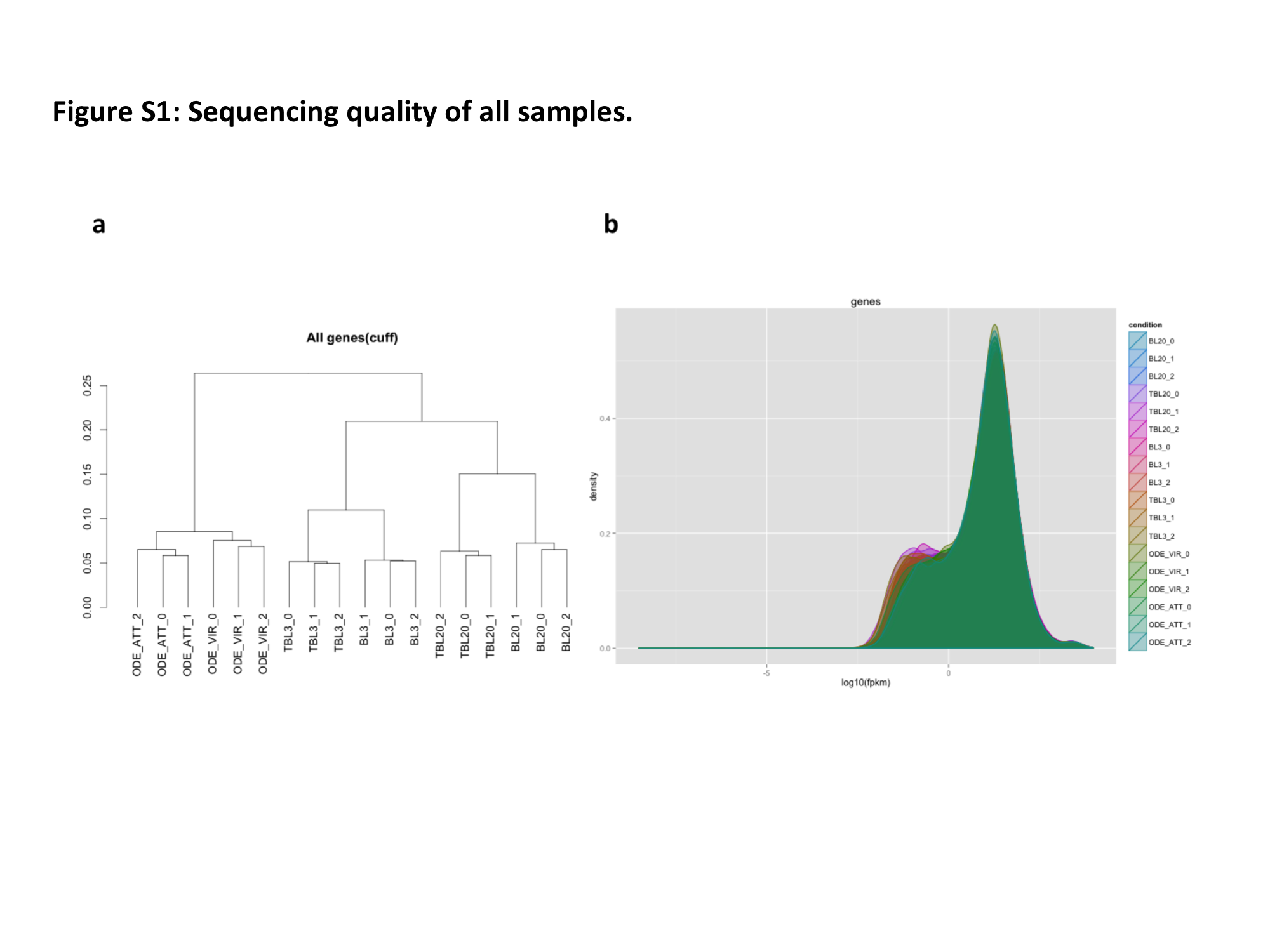

### Fig S2

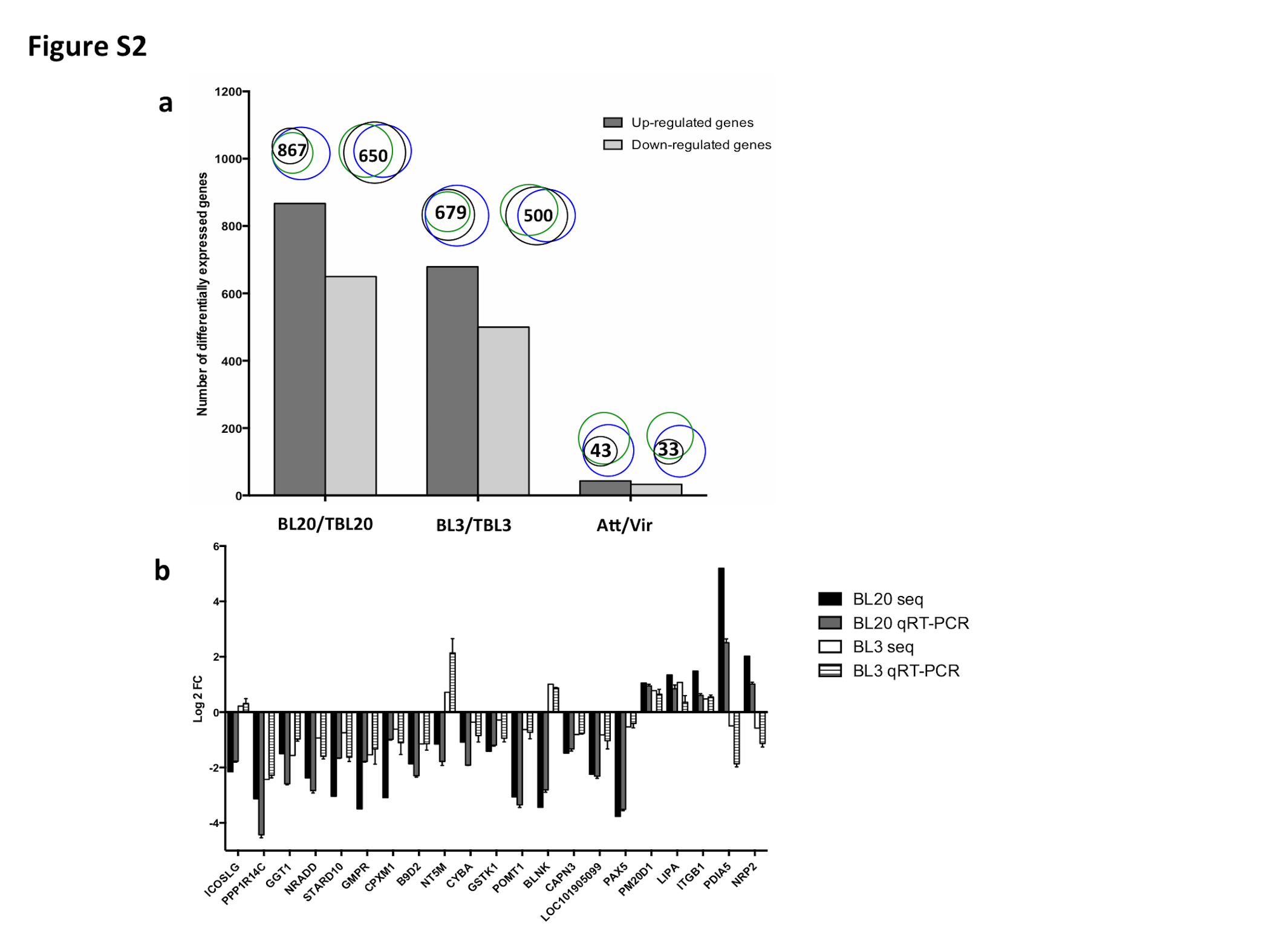

### Fig S3

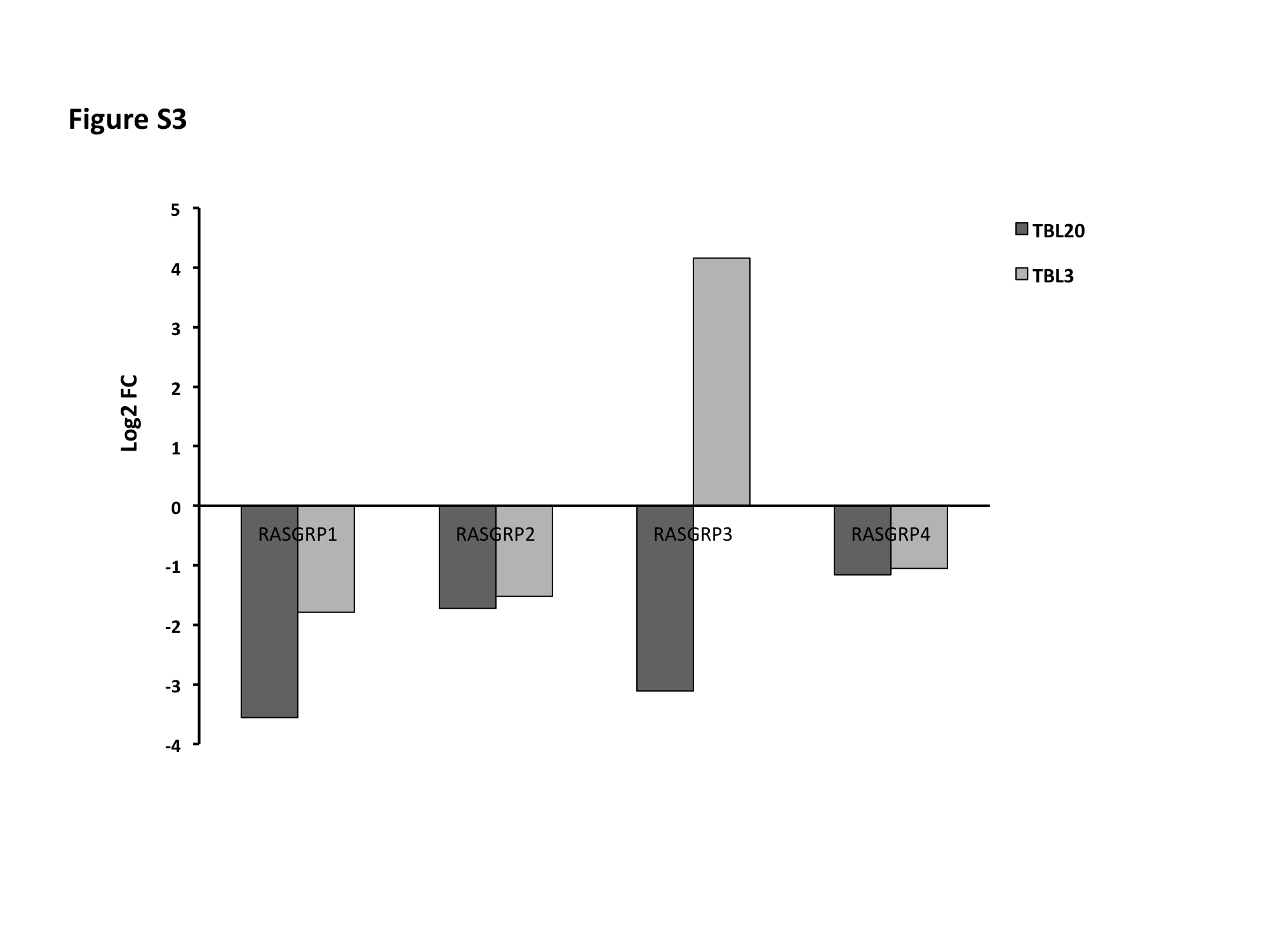

### Fig S4

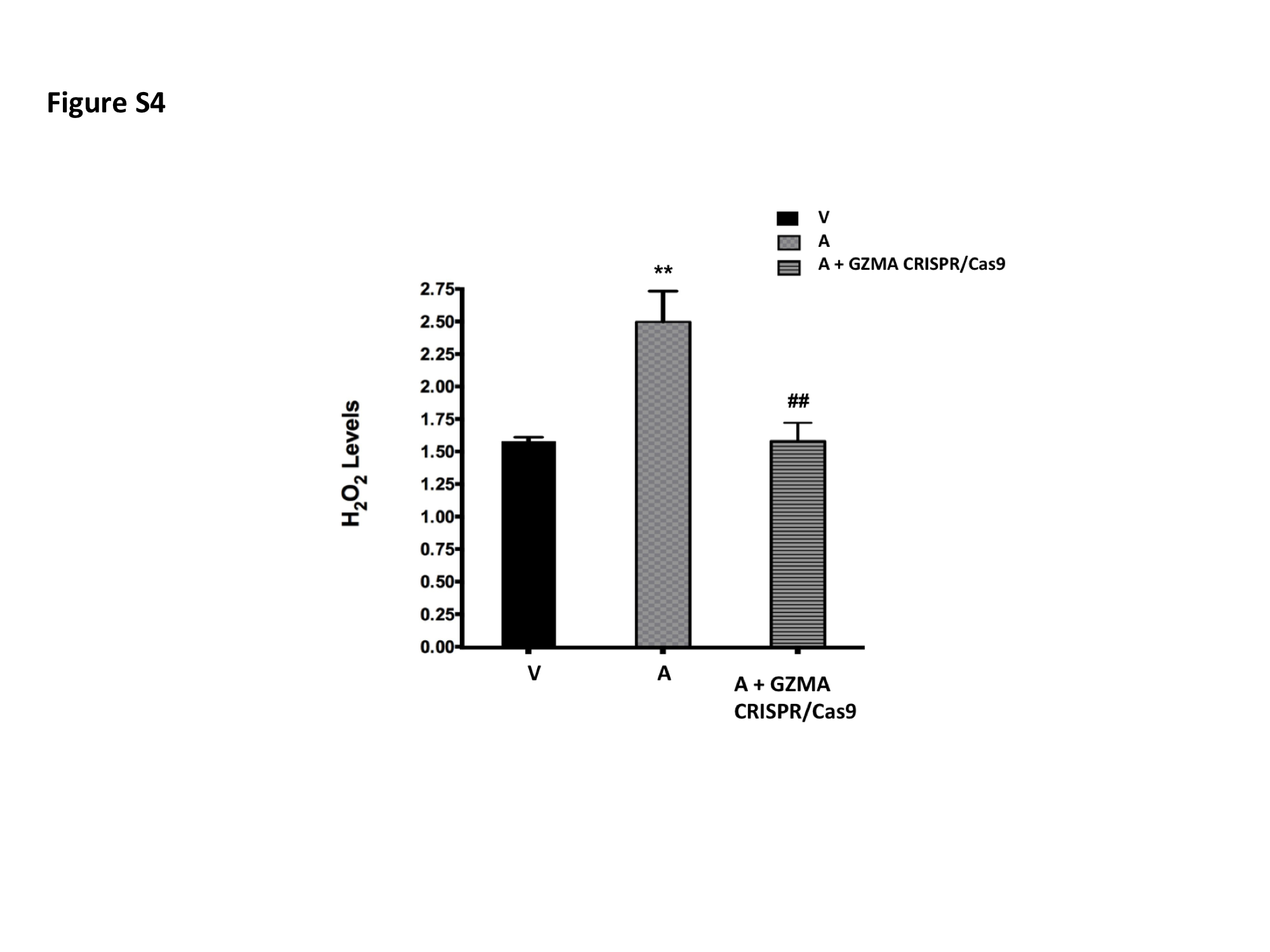

### Fig S5

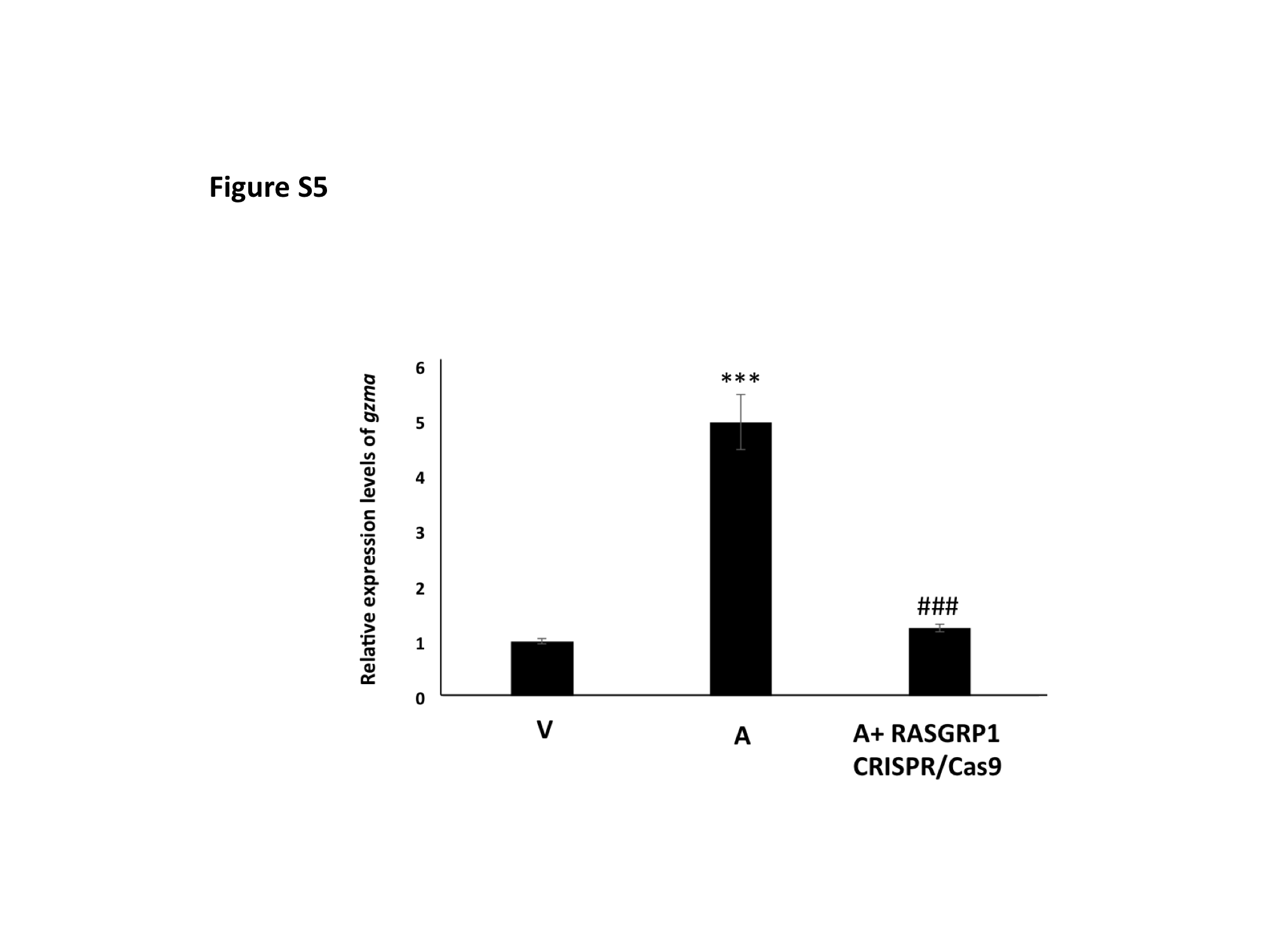

### Fig S6

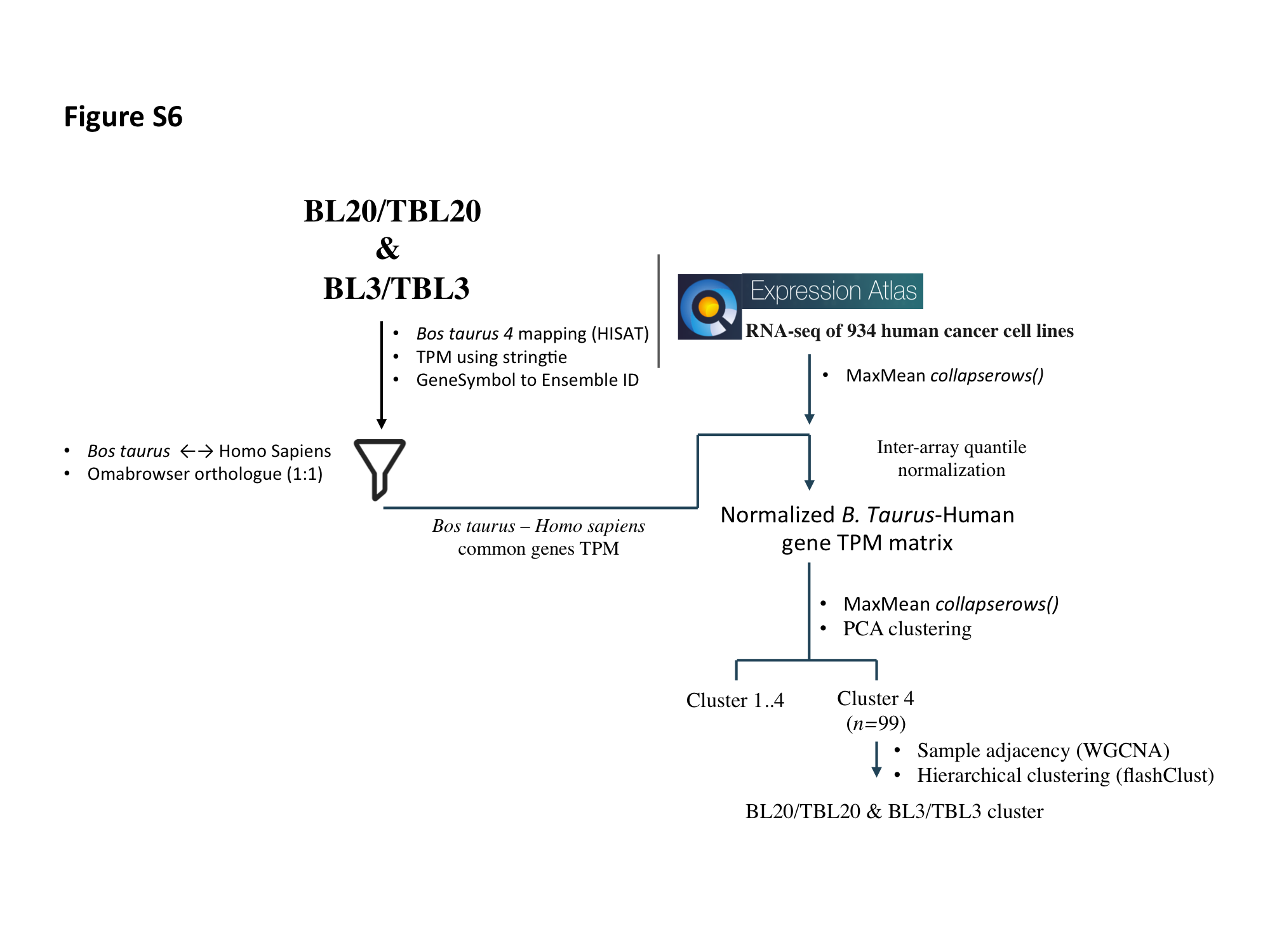

### Fig S7

#### Figure S7

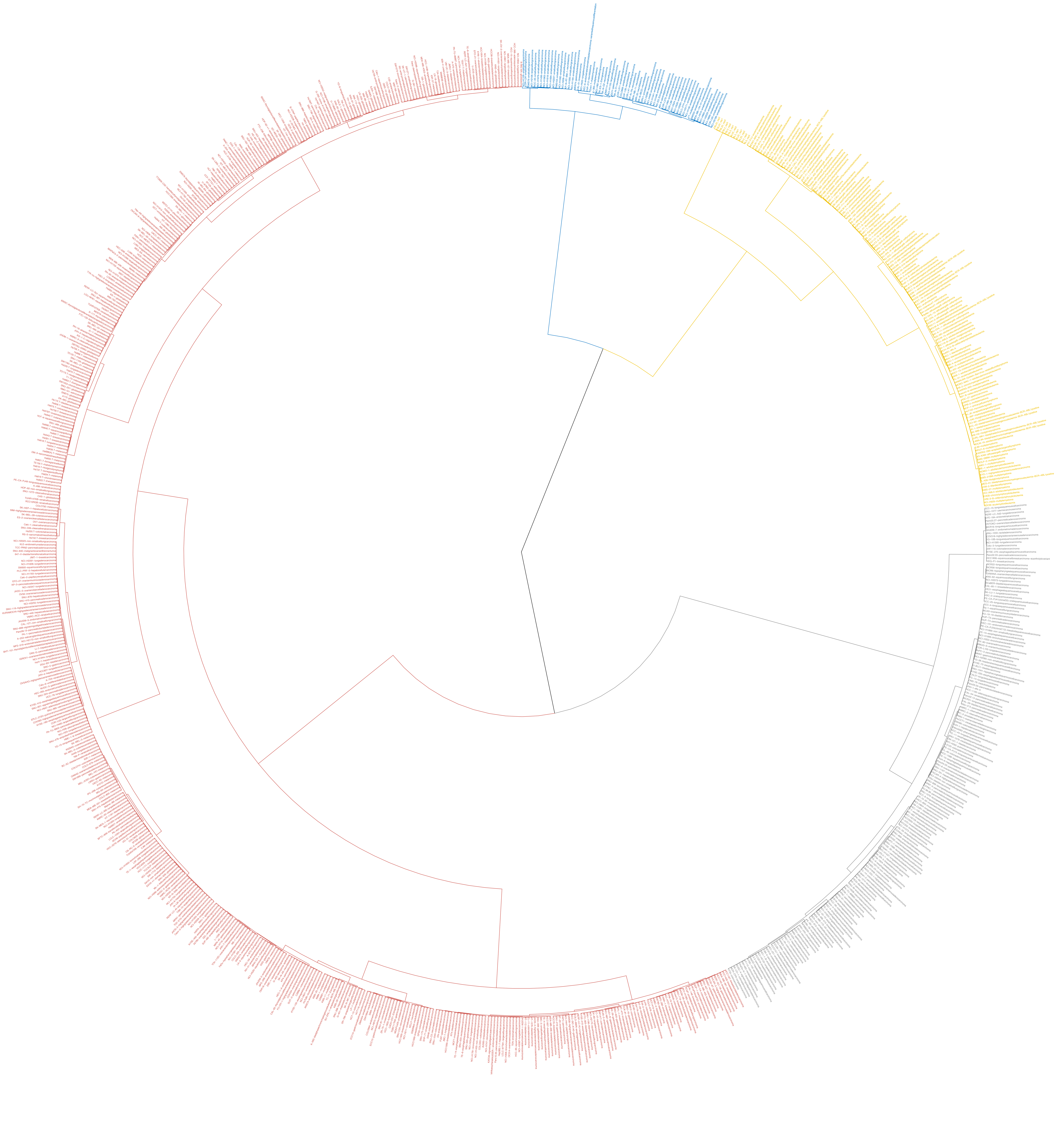
